# HAP-multitag, a PET and positive MRI contrast nanotracer for the longitudinal characterization of vascular calcifications in atherosclerosis

**DOI:** 10.1101/2020.10.20.345843

**Authors:** Juan Pellico, Irene Fernández-Barahona, Jesús Ruiz-Cabello, Lucía Gutiérrez, María J. Sánchez-Guisado, Irati Aiestarán-Zelaia, Lydia Martínez-Parra, Ignacio Rodríguez, Jacob Bentzon, Fernando Herranz

**Author notes:** These authors contributed equally (Juan Pellico, Irene Fernández-Barahona). Corresponding author at: Fernando Herranz, Instituto de Química Medica (IQM), Consejo Superior de Investigaciones Científicas (CSIC), 28006 Madrid, Spain.

## Abstract

Vascular microcalcifications are associated with atherosclerosis plaque instability and, therefore, to increased mortality. Because of this key role, several imaging probes have been developed for their *in vivo* identification. Among them [^18^F]FNa is the gold standard, showing a large uptake in the whole skeleton. Here, we push the field towards the combined anatomical and functional early characterization of atherosclerosis. For that, we have developed HAP-multitag, a bisphosphonate-functionalized ^68^Ga magnetic nanoparticle showing high affinity towards most common calcium salts present in microcalcifications, particularly hydroxyapatite. We characterized this interaction *in vitro* and *in vivo*, showing a massive uptake in the atherosclerotic lesion identified by PET and positive contrast MRI. In addition, this accumulation was found to be dependent on the calcification progression, with a maximum uptake in the microcalcification stage. These results confirmed the ability of HAP-multitag to identify vascular calcifications by PET/(*T*_1_)MRI during the vulnerable stages of the plaque progression.

## INTRODUCTION

Atherosclerosis is a complex chronic inflammatory disease of the blood vessel wall in which plaques build up inside the arteries, being the leading cause of cardiovascular diseases. A key factor in atherosclerosis development is the formation of calcified micronodules after a first inflammation step.^1^ These microcalcifications are associated with plaque rupture, leading to a cardiac event, or with plaque stabilization through the formation of macroscopic crystals (macrocalcifications) later in plaque development.^1,2^ Atherosclerosis microcalcifications are mainly composed of a mixture of hydroxyapatite (HAP), calcium oxalate monohydrate and β-tricalcium phosphate, with HAP as the major component.^3^ Due to the relevance of these microcalcifications, several imaging probes have been developed in the last years. There are two main approaches to develop tracers for *in vivo* detection of calcifications: the use of [^18^F]FNa and bisphosphonate-based tracers. [^18^F]FNa is the gold standard for PET detection of calcifications in the clinical scenario, owing to the favorable pharmacokinetic profile and lack of toxic effects.^4^ On the other hand, the [^18^F]FNa only binds to HAP, while bisphosphonate-based (BP) tracers or nanoparticles cover a broader spectrum of calcium salts recognition, relevant in several diseases.^5–8^ The mechanism of accumulation in calcifications is different for both types of tracers: in using [^18^F]FNa, ^18^F substitutes one hydroxyl group in the HAP matrix, forming fluorapatite, while when using BP-based probes, the bisphosphonate moiety coordinates with the Ca atom. When using [^18^F]FNa or BP-based tracers, one of the main drawbacks is the high uptake they show in bone, increasing the off-target signal, often complicating vasculature differentiation.^9^ If the main focus is atherosclerosis, imaging a tracer for which bone signal is minimized is highly desirable for imaging purposes. This limitation is overcome in humans and large animal models by selecting regions of interest in the imaging acquisition or post-processing steps. However, this strategy is impractical in small animal models where the vasculature is extremely small and PET resolution, even when combined with computed tomography (CT), is flawed. To the best of our knowledge, there is no reported PET radiotracer for vascular microcalcifications in atherosclerosis mouse models. Examples of microcalcification detection in mice have been exclusively described in breast cancer and chronic tuberculous models.^10–12^

A second key aspect is the imaging modality. Current probes for vascular calcification detection are mainly based on nuclear imaging techniques, particularly PET. This technique offers unparalleled sensitivity but poor spatial resolution. For this reason, PET scanners are combined with Computed Tomography (CT) and, more recently, with Magnetic Resonance Imaging (MRI) scanners providing detailed functional and anatomical information with micron resolution.^13^ The combination of PET with MRI is arguably the most convenient since it pieces together the extraordinary sensitivity of PET with the excellent resolution of MRI.^14,15^ The development of this technology is associated with the design of novel probes, providing signal in both imaging techniques. Among the different chemical compounds used to produce dual PET/MRI probes, iron oxide nanoparticles possess several advantages and one major drawback. Iron oxide nanoparticles (IONPs) are biocompatible, easy to produce and with a large variety of possible coatings to tune their bioconjugation and biodistribution.^16^ IONPs have a single drawback for this application, however, it is a major one: the typical signal they provide is T_2_-based, negative, or dark. This option complicates *in vivo* uptake identification, particularly in regions where an endogenous dark signal is present, like calcified vascular areas. This problem has drastically limited their use, especially in the clinical area, in molecular imaging or multimodal approaches. This absence has boosted the quest for IONPs, providing positive contrast in MRI, with several examples in the literature where positive contrast is achieved by tuning the core size,^17,18^ the coating size,^19^ or the core composition.^20^ Most of the time, the positive contrast is demonstrated by *in vivo* MR angiography. The dilution and sample redispersion in a large blood volume reduce the T_2_ effect, which could hamper the generation of positive contrast. However, examples of positive contrast in which IONPs accumulate in a specific tissue or organ are scarce.^20^

Here, we use bisphosphonate-based ^68^Ga-core doped iron oxide nanoparticles that we termed HAP-multitag, with several key features: they provide a simultaneous signal in PET and –positive contrast–MRI. HAP-multitag binds predominantly to HAP and other calcium salts relevant to vascular calcification, as demonstrated *in vitro* by different techniques. *In vivo*, HAP-multitag accumulation in atherosclerosis lesions can be monitored by PET and positive contrast MRI techniques. Finally, the accumulation is dependent on the progress of the lesion, which further demonstrates the ability of HAP-multitag to diagnose and longitudinally characterize, for the first time, atherosclerotic lesions by PET/(*T*_1_)MRI in mice.

## METHODS

^68^Ga (t_1/2_ = 68 min, β+ = 89% and EC = 11%) was obtained from a ^68^Ge/^68^Ga generator system (ITG Isotope Technologies Garching GmbH, Germany) in which ^68^Ge (t_1/2_ = 270 d) was attached to a column based on organic matrix generator. The ^68^Ga was eluted with 4 mL of 0.05 M hydrochloric acid. Iron (III) chloride, hydrazine monohydrate, N-(3-Dimethylaminopropyl)-N’-ethylcarbodiimide hydrochloride, N-hydroxysulfosuccinimide sodium salt and alendronate sodium salt were purchased from Sigma-Aldrich. Citric acid trisodium salt dihydrate was purchased from Acros organics. OsteoSense 680TM EX was purchased from Perkin Elmer, disposable PD10 desalting salt columns were purchased from GE Healthcare Life Sciences and Amicon® Ultra centrifugal filters from Merck Millipore.

### Synthesis of ^68^Ga-IONP-citrate

FeCl_3_ × 6 H_2_O (75 mg, 0.28 mmol), sodium citrate hydrate (80 mg, 0.27 mmol) and 1280 MBq of ^68^GaCl_3_ in HCl (0.05 M, 4 mL) were dissolved in water (5 mL) in a microwave-adapted flask, followed by addition of 1 mL hydrazine hydrate. The solution was ramped to 120°C over 54 s and held at this temperature for 10 minutes (240 W) in a Monowave 300 microwave reactor equipped with an internal temperature probe and an external IR probe (Anton Paar, GmbH, Ostfildern-Scharnhausen, Germany). The reaction mixture was then cooled to 60 °C, and the ^68^Ga-IONP-citrate product purified by passing the mixture through a PD-10 column to eliminate excess small reagents, including all unincorporated radioisotope. This purification process provided 9 mL of ^68^Ga-IONP-citrate with a total activity of 781 MBq (measured 40 minutes after starting the reaction), a radiolabeling yield of 92%.

### Synthesis of ^68^Ga-IONP-alendronate (HAP-multitag)

To 750 MBq of ^68^Ga-IONP-citrate (5 mL) were added 0.07 mmol of N-(3-dimethylaminopropyl)-N′-ethylcarbodiimide hydrochloride (EDC) and 0.075 mmol of N-hydroxysulfosuccinimide sodium salt (sulfo-NHS). The solution was stirred for 30 min at room temperature (r.t.) and then ultracentrifuged at 10,350 x g through Amicon Ultra-15 30 kDa centrifugal filters for 4 min to remove excess reagents. The retentate was resuspended in 1.5 mL HEPES buffer, pH 8, and 1 mg of alendronate sodium salt added to the solution. The mixture was maintained at r.t for 60 min with stirring. Finally, another ultrafiltration step was performed to eliminate unreacted alendronate. The retentate was resuspended in saline solution giving 195.6 MBq of HAP-multitag with a radiolabeling yield of 98%.

### Physicochemical characterization

Hydrodynamic size and polydispersity index were measured with a Zetasizer Nano ZS90 system (Malvern Instruments, UK) using folded capillary cells. For determination of morphology and mean particle size and distribution, samples were examined under a transmission electron microscope (Tecnai F30, FEI) operated at 300 KV using scanning-transmission imaging with a high angle annular dark field detector (STEM-HAADF. A drop of the nanoparticle suspension was deposited onto a holey-carbon-coated copper grid and left to evaporate at room temperature.). Mean sizes and standard deviations were calculated for approximately 50 particles.

### Titration of the Ca^2+^ salts

^68^Ga-IONP-citrate and ^68^Ga-IONP-alendronate were incubated with different concentrations (0.2 μM, 0.4 μM, 1 μM, 2 μM, 4 μM, 10 μM, 20 μM) of three different calcium salts: hydroxyapatite, calcium oxalate monohydrate and β-tricalcium phosphate. After 60 min of incubation at r.t. hydrodynamic size of the samples was measured using a Zetasizer Nano ZS90 system (Malvern Instruments, UK).

### Binding quantification by fluorescence

Alexa 647 (A647) dye (excitation λ = 649 nm;_emission λ = 666 nm)_was used to quantify binding (%) of ^68^Ga-IONP-citrate and ^68^Ga-IONP-alendronate to different calcium salts: hydroxyapatite, calcium oxalate monohydrate and β-tricalcium phosphate. To synthesize ^68^Ga-IONP-citrate-A647 and ^68^Ga-IONP-alendronate-A647: 5 mL of ^68^Ga-IONP-citrate were added to 0.07 mmol N-(3-dimethylaminopropyl)-N′-ethylcarbodiimide hydrochloride (EDC) and 0.075 mmol of N-hydroxysulfosuccinimide sodium salt (sulfo-NHS). The solution was stirred for 30 min at room temperature (r.t.) and then ultracentrifuged at 10,350 x g through Amicon Ultra-15 30 kDa centrifugal filters for 4 min to remove excess reagents. The retentate was resuspended in 1.5 mL HEPES buffer, pH 8, and 100 μg of Alexa 647 hydrazide to synthesize ^68^Ga-IONP-citrate-A647; and 100 μg of Alexa 647 hydrazide plus 1 mg of alendronate sodium salt to obtain ^68^Ga-IONP-alendronate-A647. The samples were maintained at r.T. for 60 min under vigorous stirring. Once this step was finished, samples were purified by ultrafiltration to eliminate unreacted A647 and alendronate. The retentate was resuspended in saline solution.

^68^Ga-IONP-citrate-A647 and ^68^Ga-IONP-alendronate-A647 were incubated for 60 min at room temperature with different concentrations of the calcium salts (0.2 μM, 0.4 μM, 1 μM, 2 μM, 4 μM, 10 μM, 20 μM). Posteriorly, supernatant fluorescence was measured at λ = 666 nm after 150 min centrifugation at 13,680 x g.

The degree of Ca salt binding was assessed using the following formula:

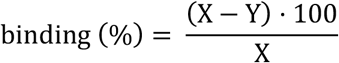

being X the initial amount of fluorescence in ^68^Ga-IONP-citrate-A647 and ^68^Ga-IONP-alendronate-A647, and Y the amount of fluorescence left in the supernatant after centrifugation.

### Animal model

Mice were housed in the specific pathogen-free facilities at the Centro Nacional de Investigaciones Cardiovasculares Carlos III, Madrid. All animal experiments conformed to EU Directive 2010/63EU and Recommendation 2007/526/EC, enforced in Spanish law under Real Decreto 53/2013. Protocol approved by Madrid regional government (PROEX16/277).

ApoE^−/−^ mice were fed with high-cholesterol diet from 8 weeks old onwards to obtain the atherosclerosis mouse model.

### PET/CT imaging

*In vivo* PET/CT imaging in mice was performed with a nanoPET/CT small-animal imaging system (Mediso Medical Imaging Systems, Budapest, Hungary). List-mode PET data acquisition commenced 90 min after injection of a bolus of 10 - 15 MBq of HAP-multitag through the tail vein and continued for 30 minutes. At the end of PET, a microCT was performed for attenuation correction and anatomic reference. The dynamic PET images were reconstructed in a 105 × 105 matrix (frame rates: 3 × 10 min, 1 × 30 min, 1 × 60 min) using a Tera-Tomo 3D iterative algorithm. Images were acquired and reconstructed with proprietary Nucline software (Mediso, Budapest, Hungary). Images were analyzed using Horos software v.3.3.6.

### Fluorescence imaging

Experiments were conducted following the standard protocol provided by the manufacturer. OsteoSense® 680EX was reconstituted by addition of 1.2 mL of PBS 1X into the vial. The mixture was gently shaken for 5 min at r.t. Then, 100 μL of the resultant solution were intravenously injected into 5 ApoE^−/−^ mice. 24 h post injection animals were sacrificed in a CO_2_ chamber, perfused with 8 mL of PBS 1X and the aortas excised. *Ex vivo* imaging of the aortas was carried out in an IVIS Imaging System 200, Xenogen® (acquisition parameters: Cy5.5 ex/em filter, high level, BIN-HR, FOV 13.3, f2, 4s).

### MRI acquisition

All experiments were performed on a 7 Tesla Bruker Biospec 70/30 USR MRI system (Bruker Biospin GmbH, Ettlingen, Germany), interfaced to an AVANCE III console. Anesthesia was induced with 3% isofluorane in 30% oxygen and maintained 1-2% isofluorane along the experiment.

A BGA12 imaging gradient (maximum gradient strength 400 mT/m) system with a 40 mm diameter quadrature volume resonator was used for MRI data acquisition. Animal were prone positioned in a customized 3D printed bed with head holder, and kept warmed with heated air pumped through an MRI compatible system interfaced to a Monitoring and gating Model 1025 (SA instruments). Temperature-control (anal) and respiration (through a respiratory pad) was registered along the experiment.

To ensure an accurate positioning, pure axial and 4-chamber view scout images were used to set up the representative aortic arch view. From these, images were acquired between the brachiocephalic artery and left common carotid artery, perpendicular to the direction of the flow in the aorta. A single 0.8 mm, 2.8×2.8 cm isotropic FOV (acquired and reconstructed with 256×256) slice was acquired using a Bruker self-gated cine gradient echo FLASH sequence using the following parameters: minimum TE 4 ms, TR 9 ms, flip angle 10, 1 average. An additional image in the same position was acquired with a fat suppression module.

### *Ex vivo* biodistribution

Biodistribution was studied with a Wizard 1470 gammacounter (Perkin Elmer). Animals were sacrificed in a CO_2_ chamber, after which blood was extracted and the animals perfused with 8 mL PBS 1X. Organs were extracted and counted in the gammacounter for 1 min each. Readings were decay corrected and presented as the percentage of injected dose per gram (%ID/g).

### Histological analysis

Excised aortas were fixed in 10% formalin for 24 hours. Tissue was dehydrated and embedded in paraffin until sectioning. Aorta sections were stained with Perl’s Prussian Blue. Images were processed and digitalized with NIS-Elements acquisition software.

## RESULTS AND DISCUSSION

### Synthesis and characterization of ^68^Ga-IONP-alendronate

We synthesized HAP-multitag (^68^Ga-IONP-alendronate) in a two-step synthetic procedure (Scheme 1). Firstly, a microwave-driven protocol rendered ^68^Ga core-doped iron oxide nanoparticles coated with citric acid (^68^Ga-IONP-citrate). This methodology, previously reported by our group, produce ^68^Ga-nanoparticles with high radiolabeling yield, high radiochemical purity and stability as well as large *r*_1_ values, ensuring a remarkable response in both PET and positive contrast MRI.^21^ Then, we coupled the bisphosphonate moiety (alendronate sodium) by EDC/sulfoNHS chemistry, following a protocol previously described.^22^

**Scheme 1.**
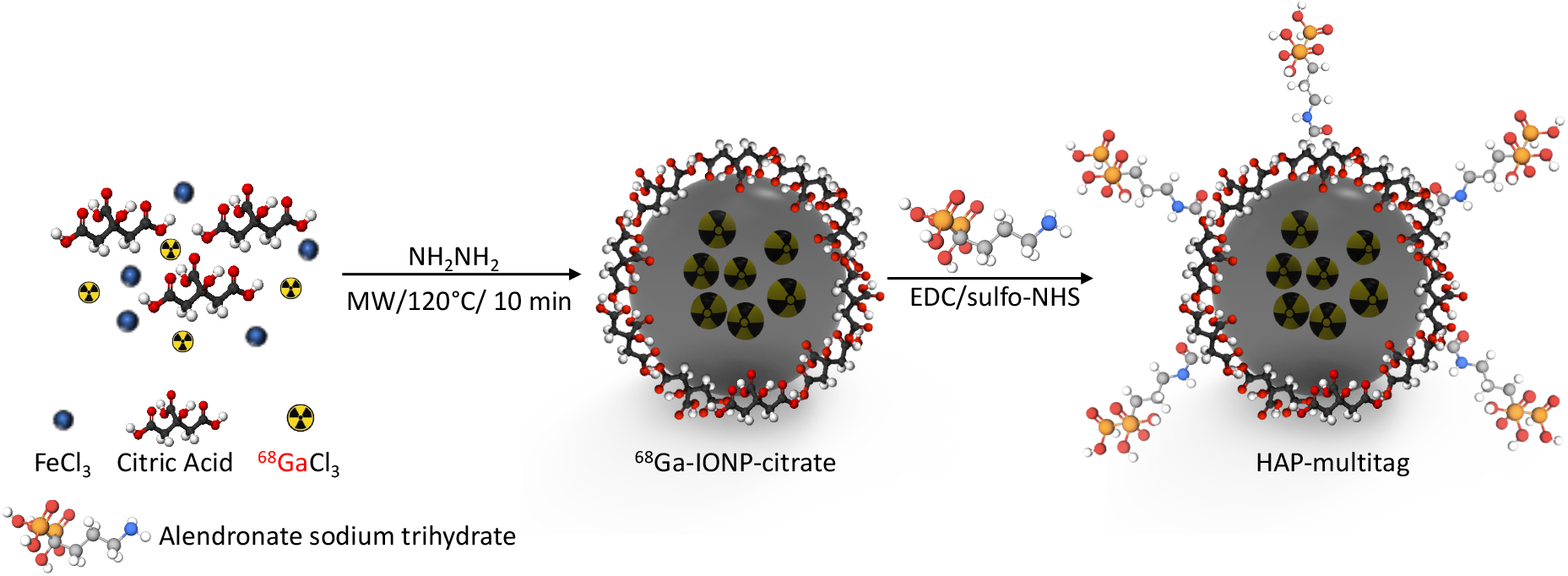
Microwave two-step synthesis of ^68^Ga-IONP-alendronate (HAP-multitag probe).

After purification by size-exclusion chromatography, we analyzed the physicochemical properties of the radiolabeled nanoparticles. Dynamic light scattering (DLS) measurements show no differences between ^68^Ga-IONP-citrate and ^68^Ga-IONP-alendronate samples (Figure 1a), indicating no aggregation after the bioconjugation step, as expected for these hydrophilic nanoparticles when using EDC and sulfoNHS as coupling agents. Z-potential measurement shows a significant reduction in the value of the superficial charge for ^68^Ga-IONP-alendronate (Figure 1b). The integration of the bisphosphonate moiety into the nanoparticle was confirmed by FTIR (Figure 1c). The ^68^Ga-IONP-alendronate spectrum shows a new area with multiple peaks of strong intensity between 1250 and 900 cm^−1^ corresponding to the vibration modes of P=O and P-OH groups and new weaker peaks between 2700 – 2200 cm^−1^ attributed to the O-H stretches of the O=P-OH groups.^23^

**Figure 1.**
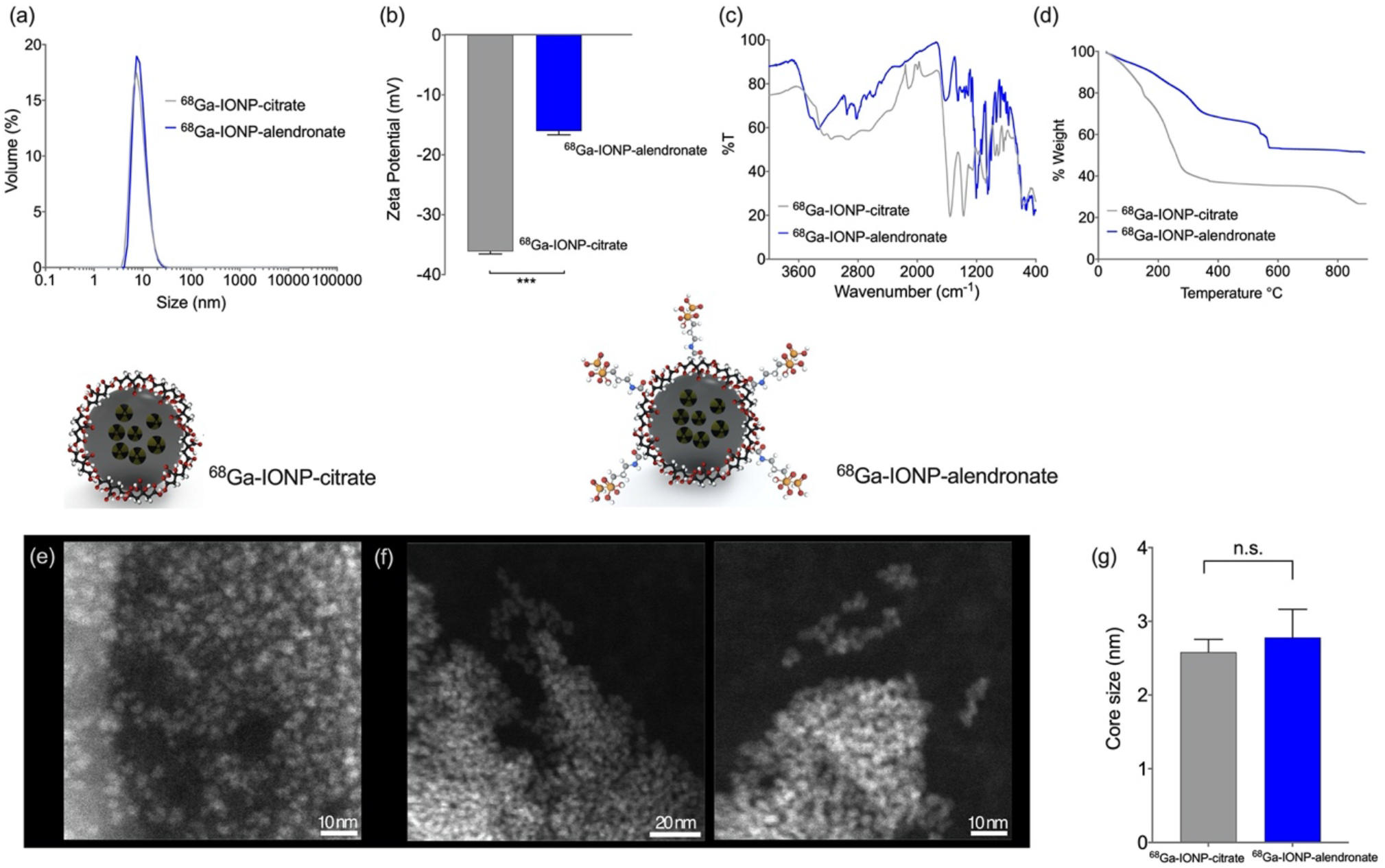
(a) DLS measurements for ^68^Ga-IONP-citrate and ^68^Ga-IONP-alendronate; (b) Z-Potential (mV) of ^68^Ga-IONP-citrate and ^68^Ga-IONP-alendronate, N=3, ***, P<0.001; (c) FTIR spectra of ^68^Ga-IONP-citrate and ^68^Ga-IONP-alendronate; (d) Thermogravimetric curves (TGA) of ^68^Ga-IONP-citrate and ^68^Ga-IONP-alendronate, e) selected STEM-HAADF image of ^68^Ga-IONP-citrate; (f) selected STEM-HAADF image of ^68^Ga-IONP-alendronate and (g) core size measured for ^68^Ga-IONP-citrate and ^68^Ga-IONP-alendronate. N=50 per sample, n.s., P>0.46.

Thermogravimetric analysis (Figure 1d) further confirms the conjugation of alendronate on the surface of the ^68^Ga-IONP-citrate, with the step between 540 – 590 °C corresponding to the covalent bond between the citric acid and the alendronate. According to TGA, the reduction in the organic coating, around 18 %, can be attributed to the loss of citrate molecules from the surface in the second reaction and purifications steps. This result (together with some exchange of citrate molecules by bisphosphonate moieties, exposing free amines) would also explain the reduction in the negative charge observed for ^68^Ga-IONP-alendronate in comparison to ^68^Ga-IONP-citrate. Finally, we studied ^68^Ga-IONP-citrate and ^68^Ga-IONP-alendronate by electron microscopy. Since these nanoparticles consist of an extremely-small iron oxide core and a large organic coating, electron microscopy images are not easily obtained. Using Scanning Transmission Electron Microscopy High-angle annular dark-field imaging (STEM-HAADF), it is possible to observe the small iron oxide cores without apparent aggregation for ^68^Ga-IONP-citrate (Figure 1e) and ^68^Ga-IONP-alendronate (Figure 1f). Analysis of the core sizes shows similar sizes for both nanoparticles, 2.6 ± 0.3 nm for ^68^Ga-IONP-citrate, and 2.8 ± 0.7 nm for ^68^Ga-IONP-alendronate.

### Qualitative assessment of the binding between ^68^Ga-IONP-alendronate and calcium salts

To assess the interaction between ^68^Ga-IONP-alendronate and calcium salts, we chose those normally present in the microcalcifications structure, i.e. hydroxyapatite (HAP), calcium oxalate monohydrate and β-tricalcium phosphate. First, we used STEM-HAADF to analyze this interaction. Figure 2a and 2c shows the different behavior of both nanoparticles when incubated with HAP. In the case of ^68^Ga-IONP-alendronate (Figure 2a), it is possible to observe the large HAP particles surrounded by the much smaller, ^68^Ga-IONP-alendronate nanoparticles, indicating their affinity towards the salt. This was further confirmed by EDX analysis, where peaks corresponding to Fe, Ca, and P where found (Figure S1). On the contrary, for ^68^Ga-IONP-citrate (Figure 2c), the HAP particles are almost free of iron oxide nanoparticles. As a qualitative technique, we used DLS, by measuring the hydrodynamic size of the nanoparticles with increasing amounts of the calcium salts, it is possible to assess whether they are interacting or not.^24,25^ We incubated the ^68^Ga-IONP-alendronate with the aforementioned salts, and measured their hydrodynamic size (Figure 2b), and similarly with ^68^Ga-IONP-citrate (Figure 2d). The Z-average value clearly shows the aggregation of ^68^Ga-IONP-alendronate as the concentration of each salt increases. This is particularly true for HAP, which shows a very large hydrodynamic size value, around 2000 nm, for the highest concentration of the calcium salt. Similarly, the interaction with the other two salts, Ca_3_(PO_4_)_2_ and Ca(COO)_2_, is clearly reflected in the aggregation of the nanoparticles. As a control, we performed the same titrations but using ^68^Ga-IONP-citrate. In this case (Figure 2d) there is no aggregation when using same calcium salts, as reflected in the constant value of hydrodynamic size (Figure 2d has the same scale in the Y axis than graph in Figure 2b for better comparison). In fact, the size measured for the highest concentration of HAP with ^68^Ga-IONP-citrate reflects the presence of the HAP nanoparticles (with a size around 300 nm) rather than an interaction between the calcium salt and the citrate nanoparticles. This result is demonstrated in Figure 2d and confirmed by the quantitative analysis (see below).

**Figure 2.**
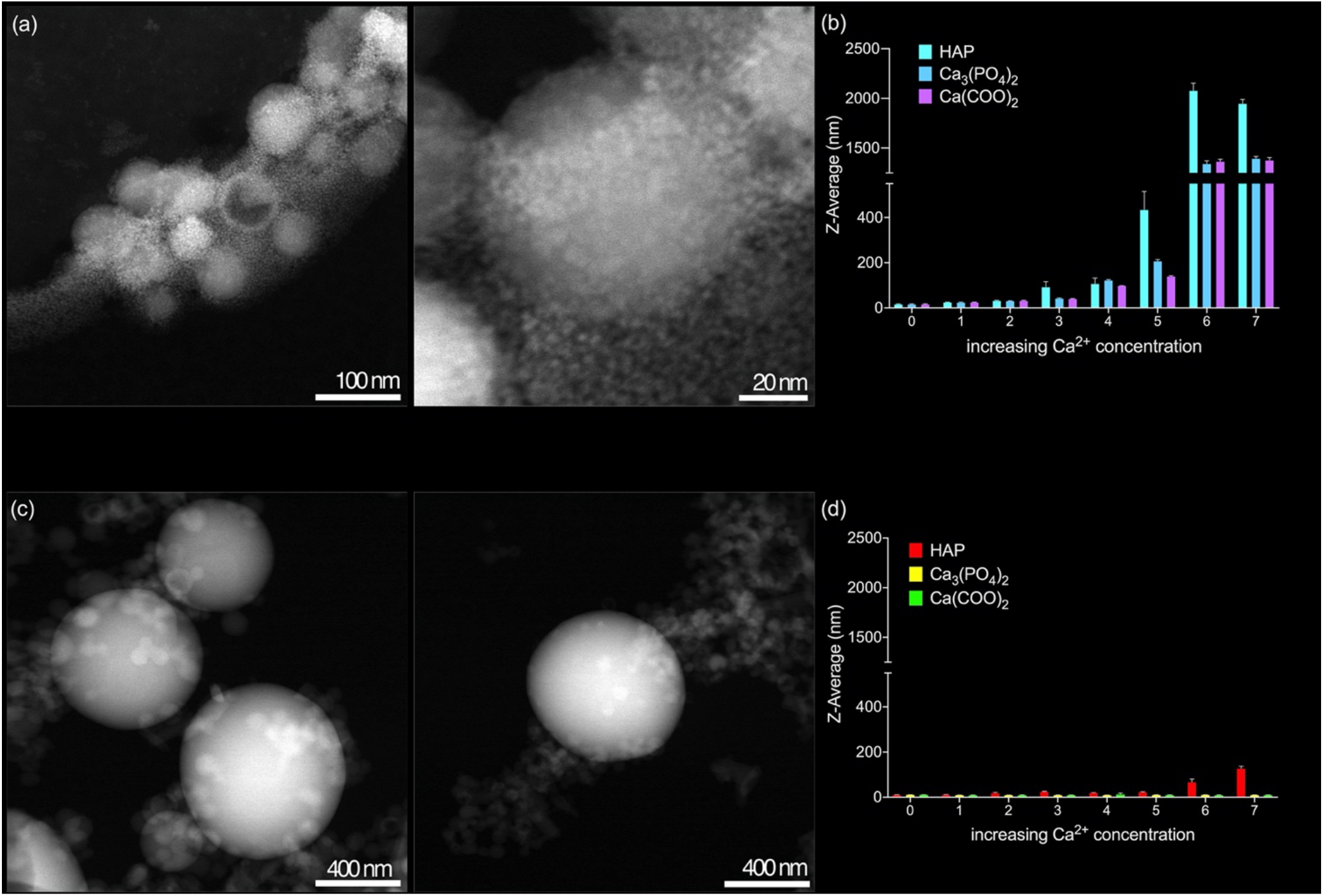
(a) STEM-HAADF images for the combination of ^68^Ga-IONP-alendronate with HAP; (b) Change in hydrodynamic size for ^68^Ga-IONP-alendronate upon the increase in the concentration of HAP, β-tricalcium phosphate and calcium oxalate monohydrate; (c) STEM-HAADF images for the combination of ^68^Ga-IONP-citrate with HAP and (d) Change in hydrodynamic size for ^68^Ga-IONP-citrate upon the increase in HAP concentration, β-tricalcium phosphate, and calcium oxalate monohydrate (scale in Y axis equal to (b) for comparative purposes).

### Quantitative assessment of the binding between ^68^Ga-IONP-alendronate and calcium salts

Next, we quantitively assessed the interaction between ^68^Ga-IONP-alendronate and ^68^Ga-IONP-citrate with the three calcium salts often present in vascular calcifications: hydroxyapatite (HAP), Ca_3_(PO_4_)_2_, and Ca(COO)_2_ (Figure 3). For this, we covalently attached to the surface of the nanotracers a fluorescent dye (Alexa 647). The nanotracers were mixed with the different salts, purified and the supernatant fluorescence quantified at each point (see methods).

**Figure 3.**
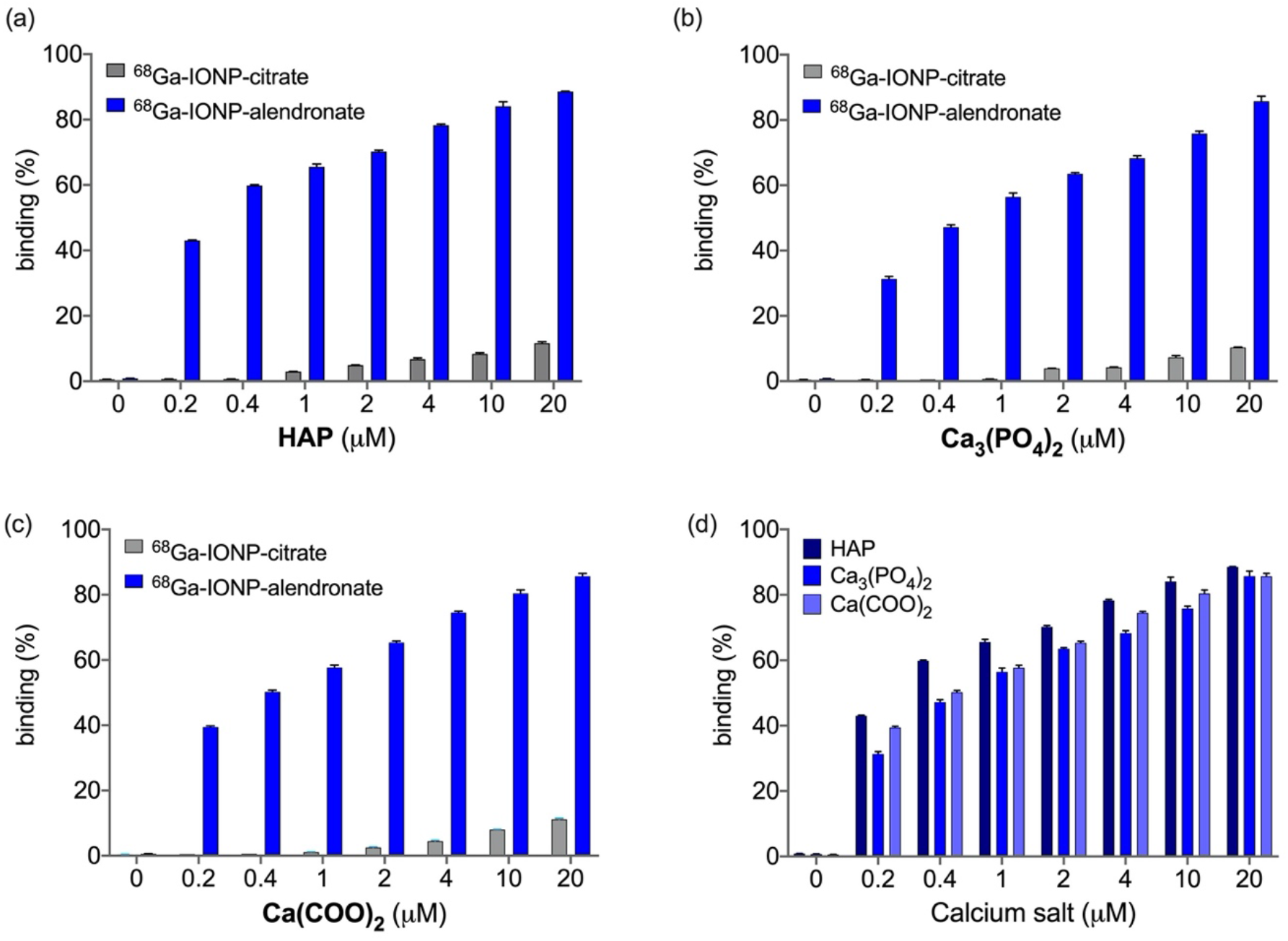
Percentage of binding between ^68^Ga-IONP-citrate and ^68^Ga-IONP-alendronate and (a) HAP; (b) Ca_3_(PO_4_)_2_ and (c) Ca(COO)_2_; (d) comparison of the binding between ^68^Ga-IONP-alendronate and the three calcium salts.

The results from these titrations confirm several aspects: First, in agreement with the DLS data, the interaction between ^68^Ga-IONP-citrate and the different salts is negligible, a mere 11 % for the largest HAP concentration (20 μM). On the contrary, titrations with ^68^Ga-IONP-alendronate clearly show a strong interaction, explaining the large aggregation observed in DLS and electron microscopy. For example, for a low concentration of HAP of 0.2 μM, the percentage of binding is already 43 %; almost half of the nanotracer sample has bound the salt at that concentration (Figure 3a). Similar results are observed for the other salts (Figure 3b,c): large binding of the alendronate nanotracer without appreciable interaction of the citrate nanotracer. Finally, a similar profile was observed for the interaction between ^68^Ga-IONP-alendronate and all the salts (Figure 3d), with a slightly stronger interaction with HAP, confirming this nanotracer presents a broad spectrum of interactions with calcium salts, not just limited to HAP, as is the case for [^18^F]FNa.

### *Ex vivo* application of ^68^Ga-IONP-alendronate; HAP-multitag

After characterizing the *in vitro* interaction between ^68^Ga-IONP-alendronate and the selected calcium salts, we tested its performance to diagnose atherosclerosis. First, *ex vivo* experiments were conducted to evaluate whether the nanotracer has any affinity towards atherosclerotic plaques in mice. Atherosclerotic ApoE^−/−^ mice were selected as a disease model. The development of hypercholesterolemia triggering aortic, carotid, and pulmonary artery lesions throughout ApoE^−/−^ mice aging is well-established.^26^ A longitudinal study was carried out in mice between 12 and 26 weeks of age. In addition, mice were fed with high-cholesterol diet from 8 weeks old onwards to accelerate atherosclerosis progression.^27^ A complete biodistribution study was performed in a gammacounter after *i.v*. injection of ^68^Ga-IONP-alendronate in ApoE^−/−^ mice. We studied four different groups: 12 weeks old and fed 4 weeks with a high-fat cholesterol diet (HFD) (Group A), 16 weeks old and 8 weeks HFD (Group B), 24 weeks old and 16 weeks HFD (Group C) and 26 weeks old and 18 weeks HFD (Group D) (Figure 4a). Main organs, blood and perfused aortas were evaluated in 5 mice of each group. Uptake values, calculated as the percentage of injected dose per gram of tissue, showed the liver and spleen as the organs with the highest accumulation. This is an expected result since biodistribution and clearance studies of iron oxide nanoparticles have demonstrated liver and spleen as the main uptake organs.^28^

**Figure 4.**
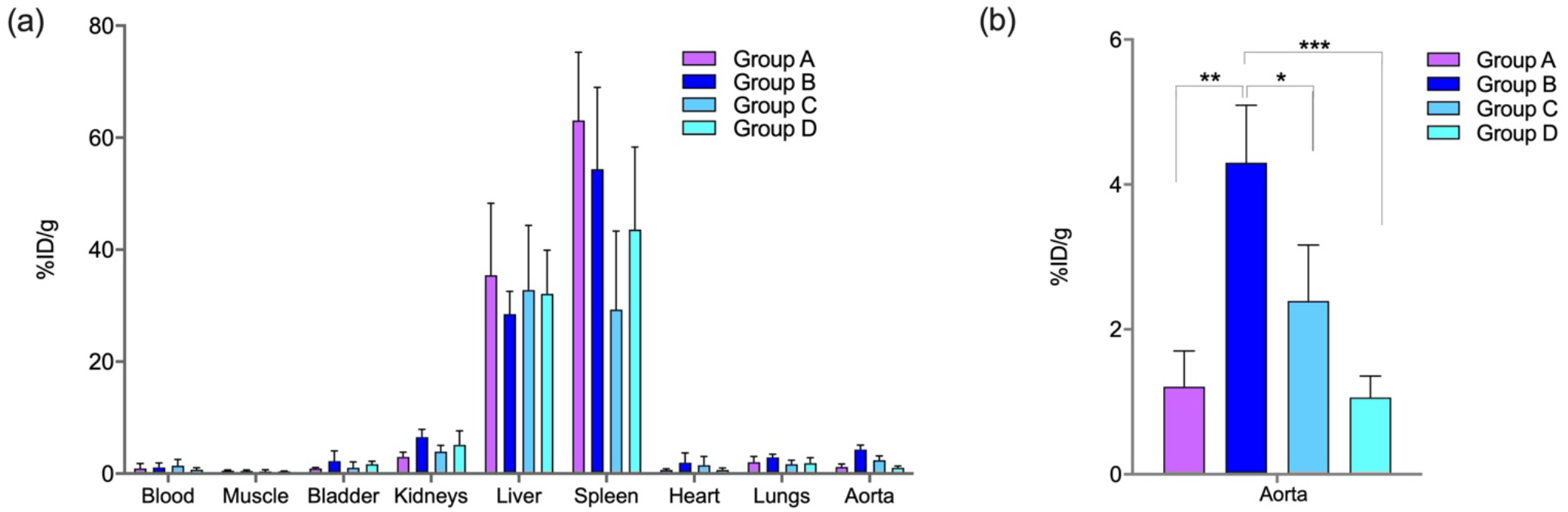
(a) Distribution of HAP-multitag measured in a gammacounter expressed as the percentage injected dose per gram (%ID/g) in ApoE^−/−^mice (N = 5) of Groups A-D; (b) Aorta uptake of HAP-multitag showing significant differences between mice groups. *P< 0.05, **P<0.01, ***P<0.001, one-way ANOVA; error bars indicate s.d., N = 5.

Aortas show an important uptake with significant differences following mice aging, hence atherosclerosis progression (Figure 4b). Negligible blood circulation of the nanoparticles (< 1.5 %ID/g) and the aortas’ perfusion, prior to the measurement, ensure that the signal measured in the aorta is due to nanoparticle uptake. Figure 3b shows the uptake of the nanoparticles in the aorta depending on the mice’s age. The HAP-multitag uptake is similar for youngest and oldest mice, with a maximum for 16 weeks old mice. This observation may have important consequences for atherosclerosis characterization. First, this profile seems to follow the calcification process: initially the amount of microcalcifications is too low to show a significant uptake —at 12 weeks— then, as more microcalcifications accumulate, an increase in the nanotracer uptake is observed —at 16 weeks— finally, the growth of the calcified deposits, and the concurrent reduction of the active surface, translates in a reduction of the nanotracer uptake, a process well-known for other tracers.^4^ Secondly, the maximum uptake for HAP-multitag the earliest reported, allowing for very early diagnosis of atherosclerosis. For comparison, the maximum uptake for [^18^F]FNa is reported in ApoE^−/−^ mice at 30 weeks old.^29^

### *In vivo* multimodal imaging of atherosclerosis with HAP-multitag

#### PET/CT imaging

Encouraged by our *ex vivo* results, we tested the ability of HAP-multitag probe to diagnose atherosclerosis by *in vivo* imaging, first using PET/CT. HAP-multitag was intravenously injected in Group B ApoE^−/−^ mice, and images recorded 90 min post injection. Spots of nanotracer uptake are observed in the aortic arch and the aorta (Figure 5a and Figure S2). PET/CT images were also acquired in Group D ApoE^−/−^ mice to determine the capacity of HAP-multitag to distinguish between micro- and macrocalcifications. Contrary to what we see in young mice, these mice showed negligible uptake in the specific regions of interest (Figure 5b), agreeing with the *ex vivo* uptake we have previously shown. Compared to[^18^F]FNa and other BP-based tracers, bone uptake is negligible (Figure 5a and Figure 5b). Nanotracer uptake was confirmed by *ex vivo* PET imaging of excised aortas after *in vivo* experiments. Signaling spots are clearly identified throughout the aorta, predominantly in the aortic arch and the renal bifurcation (Figure 5c and S2). To confirm whether the uptake is related to vascular calcifications, *ex vivo* fluorescence images were acquired using OsteoSense®. This is a commercial dye, showing fluorescence in the near infrared region, which includes a bisphosphonate moiety and it is gold-standard for *ex vivo* microcalcification detection by fluorescence techniques.^30,31^ Following the manufacturer instructions, *ex vivo* fluorescence imaging was conducted 24 h post *i.v*. injection of Osteosense® in group B ApoE^−/−^ mice (n = 5, Figure S4). Comparing the *ex vivo* PET signal (Figure 5c) with fluorescence signal from OsteoSense® (Figure 5d), there is a perfect match between the different spots showing uptake of the probes, confirming the presence of microcalcifications in the sites where there is a clear uptake of HAP-multitag. Finally, histological images show the presence of iron oxide nanoparticles, from HAP-multitag, (blue spots due to Perls’ Prussian Blue staining) in Group B ApoE^−/−^ mice but not in Group D (Figure 5e and Figure S3).

**Figure 5.**
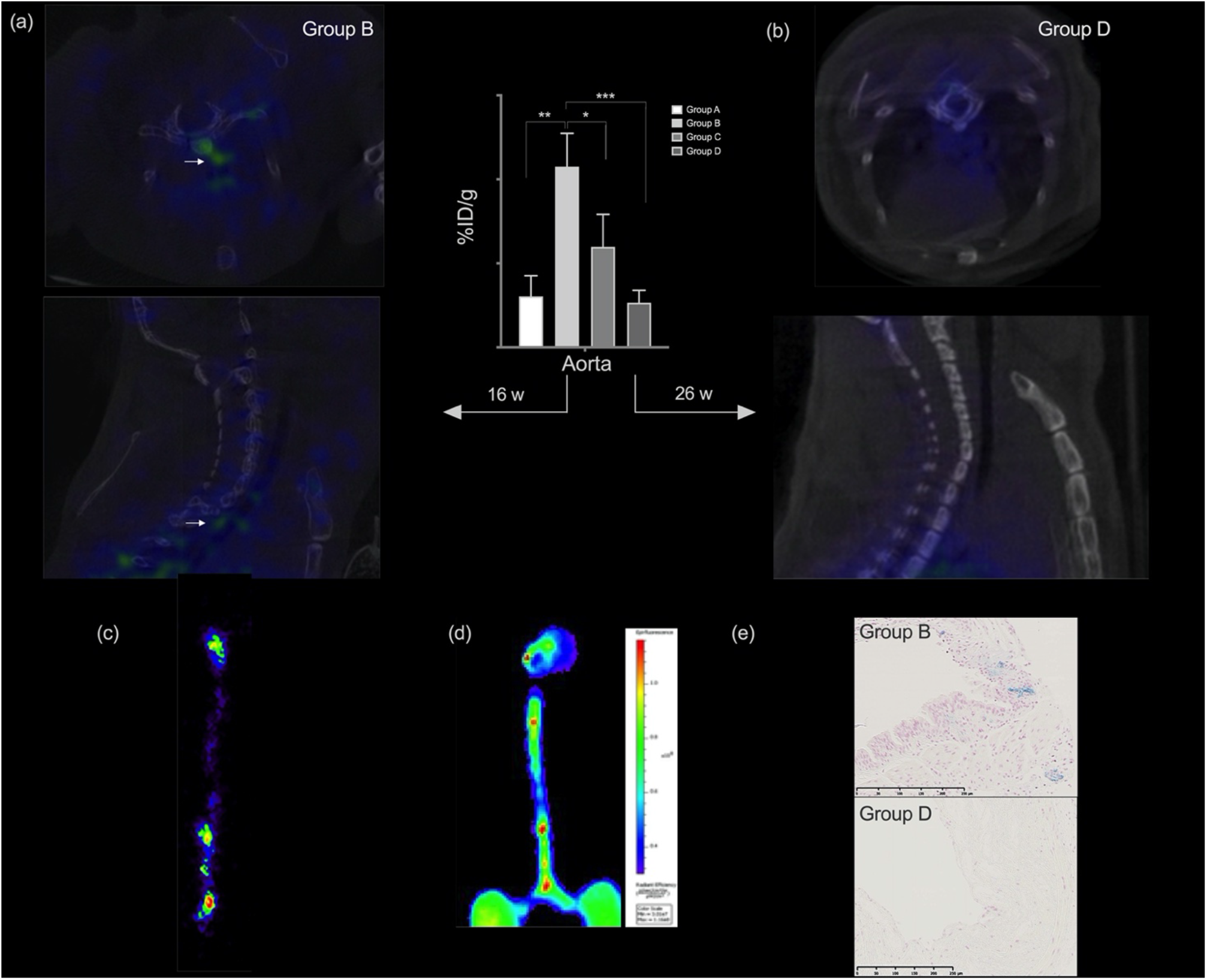
a) PET/CT imaging of a Group B ApoE^−/−^mouse 90 min post *i.v*. injection with HAP-multitag; b) PET/CT imaging of a Group D ApoE^−/−^mouse 90 min post *i.v*. injection with HAP-multitag. Central graph corresponds to Figure 4b, included here to compare uptake with images; c) *Ex vivo* PET imaging of a Group B ApoE^−/−^ mouse aorta 90 min post *i.v*. injection with HAP-multitag; d) *Ex vivo* fluorescence imaging of a Group B ApoE^−/−^ mouse aorta 24 h post *i.v*. injection of OsteoSense® 680EX; e) Perls’ Prussian blue staining of aorta sections from Group B ApoE^−/−^ mouse (top row) and Group D ApoE^−/−^ mouse (bottom row), both injected with HAP-multitag (scale bar is 250 μm). Histology of the aortas stained with Perls’ Prussian Blue confirmed the presence of iron oxide nanoparticles (blue spots, Figure 5e, top row) for Group B ApoE^−/−^ mice as well as their absence in Group D ApoE^−/−^mice (Figure 5e, bottom row).

#### Magnetic resonance imaging

Finally, the HAP-multitag performance, as positive contrast tracer in MRI, was evaluated. First, we tested the relaxometric values for HAP-multitag. At 1.5 T, the *r*_1_ value was 10.9 ± 0.1 mM^−1^s^−1^ while the *r*_2_ value was 22.0 ± 0.4 mM^−1^s^−1^. This results in a *r*_2_/*r*_1_ ratio of 1.98 ± 0.05. As extensively revised, nanoparticles with *r*_2_/*r*_1_ ratios below 4 have a high capability to provide positive MRI contrast.^17,19^ Therefore, the low ratio in HAP-multitag ensures their performance as a positive contrast agent.

*In vivo* imaging using non-functionalized ^68^Ga-IONP-citrate was conducted in Group A, Group B and Group D ApoE^−/−^ mice with no significant contrast enhancement observed in the aortic arch of these groups (Figure 6d-f and Figure S5). Then, MRI was carried out using HAP-multitag as the tracer. In this case, no significant contrast in Group A ApoE^−/−^ and Group D ApoE^−/−^ mice was observed (Figure 6a and 6c). However, the positive contrast was unambiguously appreciated 90 min. after *i.v* injection of HAP-multitag in the Group B ApoE^−/−^ mice (Figure 6b and Figure S6). Semi-quantitative analysis of the images confirmed these results. For that, ten different regions of interest (ROIs) were selected in the muscle (used as reference) and the aorta in 3 different animals per group. No significant differences were obtained using ^68^Ga-IONP-citrate, in all animals, and HAP-multitag for Group A ApoE^−/−^ and Group D ApoE^−/−^ mice (Figure 6d-f). In agreement with the *in vivo* imaging results, significant intensity differences were found for Group B ApoE^−/−^ mice. These results show the ability of HAP-multitag to generate positive contrast in MRI in a manner relevant to the calcification stage of the aorta.

**Figure 6.**
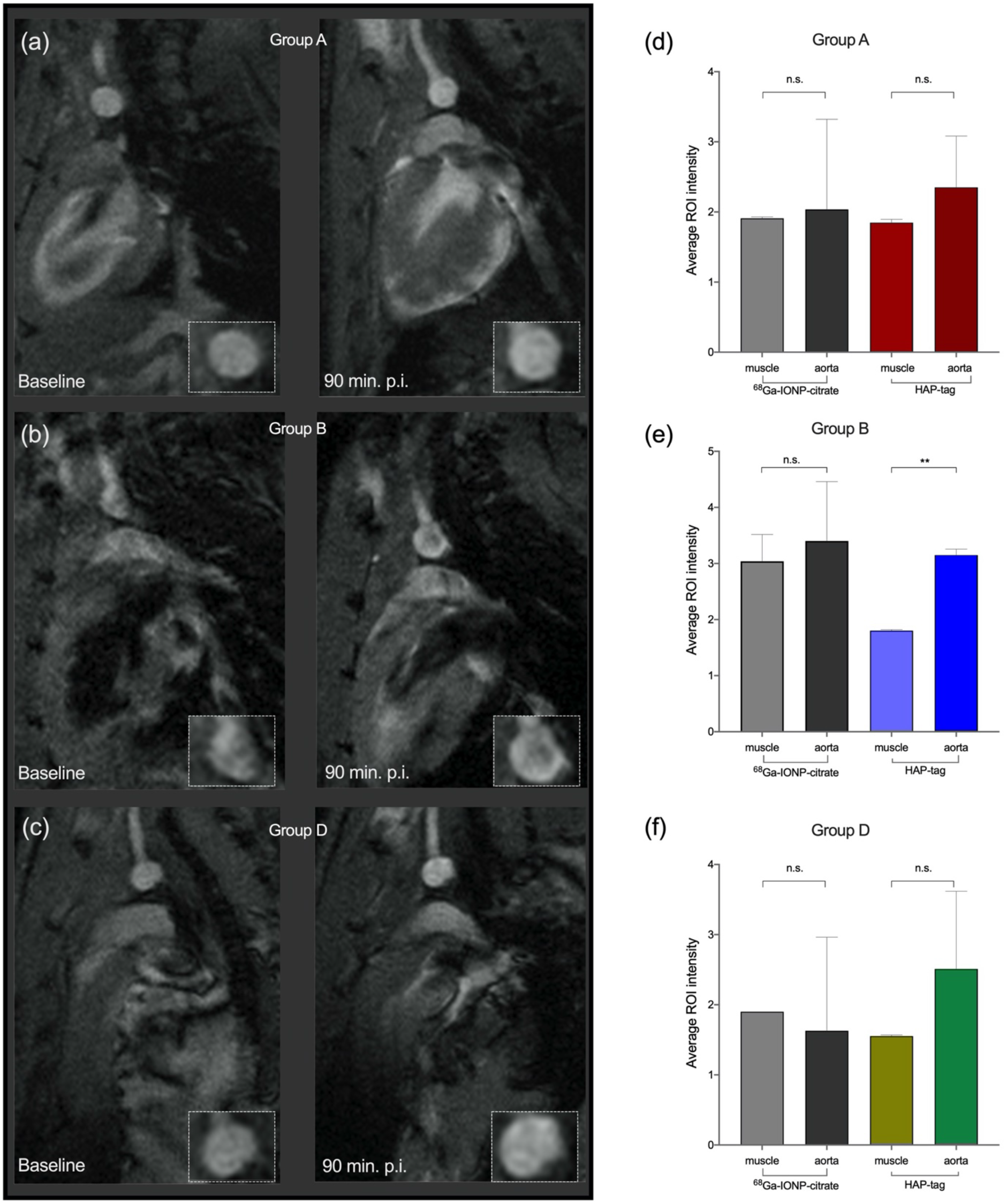
*T*_1_-weighted MRI before (baseline) and 90 min after *i.v.* injection of HAP-multitag for a) Group A ApoE^−/−^, b) Group B ApoE^−/−^ and c) Group D ApoE^−/−^; Average ROI intensity (n = 10) in muscle and aorta 90 min after *i.v.* injection of ^68^Ga-IONP-citrate or HAP-multitag in ApoE^−/−^ mice (n = 3) for d) Group A ApoE^−/−^; e) Group B ApoE^−/−^ and f) Group D ApoE^−/−^. *P< 0.05, **P<0.01, ***P<0.001, one-way ANOVA; error bars indicate s.d., N = 5.

## CONCLUSIONS

The development of PET/MRI as a powerful molecular imaging technique requires the development of imaging probes capable of providing simultaneous signal in both modalities. On this sense iron oxide nanoparticles are the perfect candidate due to their tailored synthesis, biofunctionalization and biocompatibility. They are perfect for the purpose, with the exception of one key aspect; the typical negative contrast they provide. Here, we show that it is possible to combine PET signal and positive contrast using iron oxide nanoparticles. The *in vitro* affinity of HAP-tag for calcium salts translates into an *in vivo* uptake that depends on mice age —therefore, in the calcification stage. We show how the targeted accumulation of these nanoparticles translates into easily identifiable PET and —bright signal— MRI, beyond the magnetic resonance angiography typically performed with other iron oxide nanoparticles. Using our nanotracer, HAP-multitag, it is possible to get an early characterization of atherosclerotic plaques, in mice just 16 weeks old, whose uptake permits the longitudinal characterization of microcalcifications.

## Acknowledgements

This work was supported by the Spanish Ministry of Science, grants number (SAF2016-79593-P, RED2018-102469-T and PID2019-104059RB-I00), and from the Gobierno Vasco, Dpto. Industria, Innovación, Comercio y Turismo under the ELKARTEK Program (Grant No. KK-2019/bmG19). JR-C received funding from the BBVA Foundation (Ayudas a Equipos de investigación científica Biomedicina 2018).

The CNIC is supported by the MICINN and the Pro-CNIC Foundation, and is a Severo Ochoa Center of Excellence (MICINN award SEV-2015-0505). CIC biomaGUNE is supported by the Maria de Maeztu Units of Excellence Program from the Spanish State Research Agency – Grant No. MDM-2017-0720

