## Supporting information for "HAP-multitag, a PET and positive MRI contrast nanotracer for the longitudinal characterization of vascular calcifications in atherosclerosis"

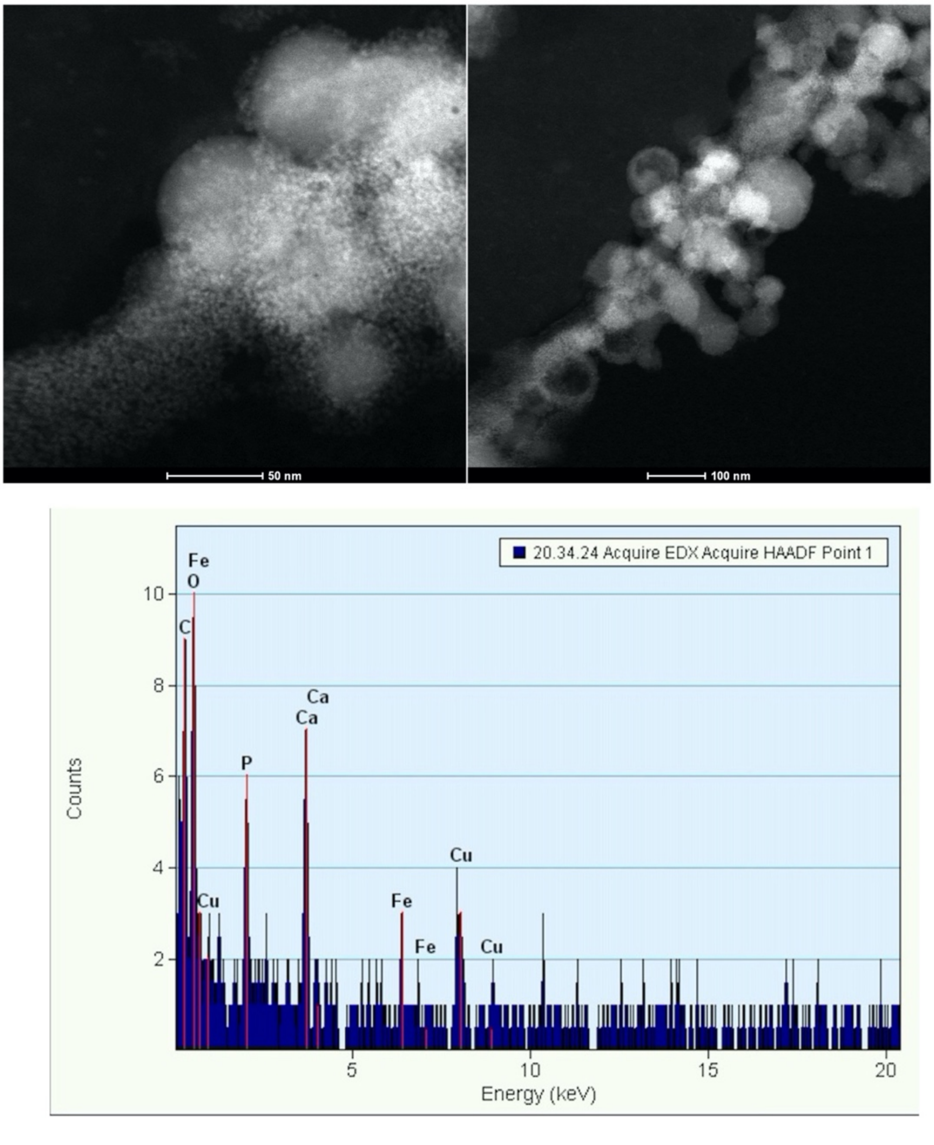


**Figure S1.** STEM-HAADF images for the combination of $^{68}$Ga-IONP-alendronate with HAP and EDX spectrum for this image.

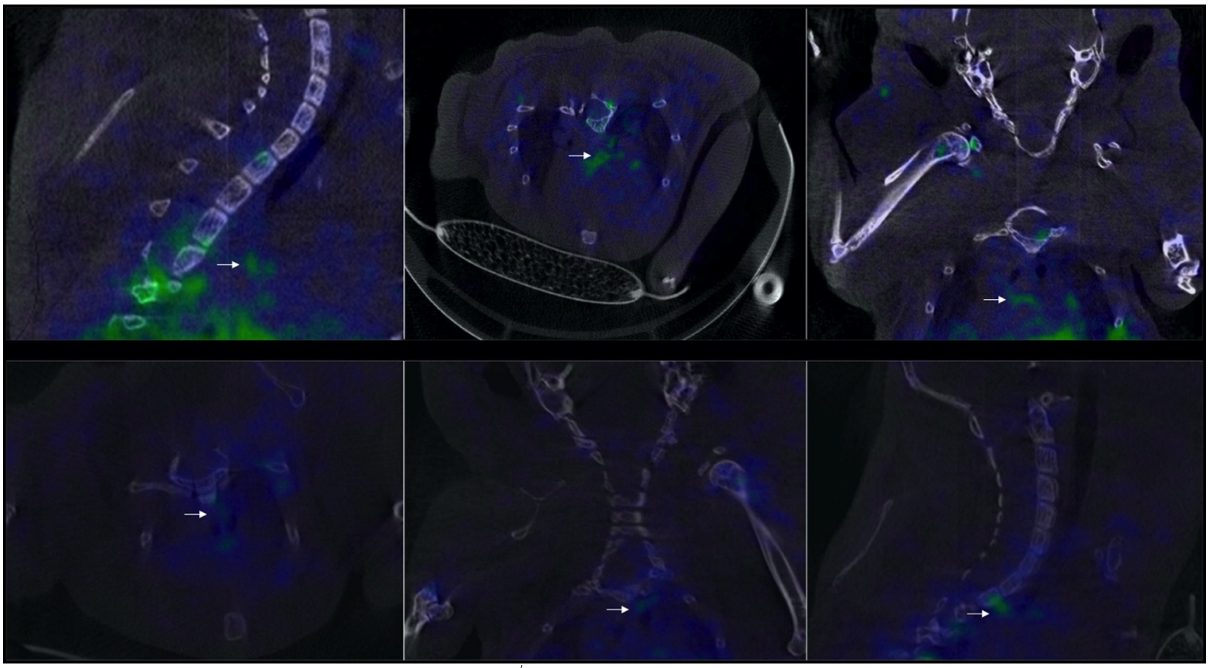

**Figure S2.** PET/CT imaging of a Group B ApoE$^{-/-}$ mice 90 min post *i.v.* injection with HAP-multitag

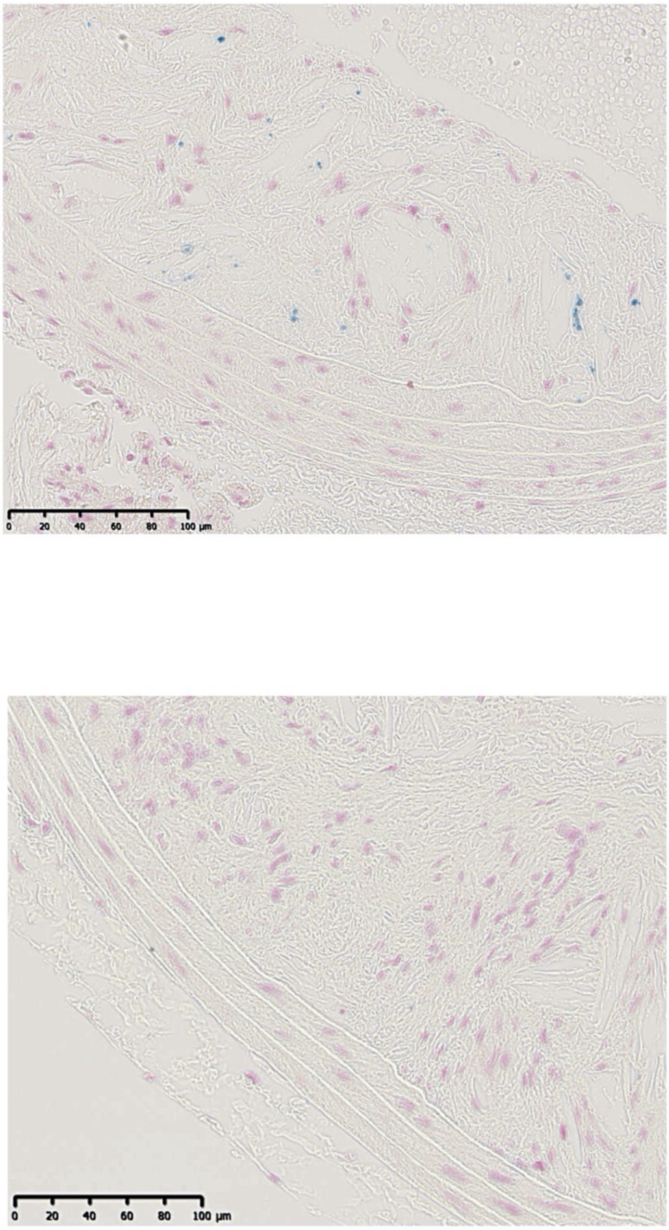


**Figure S3.** Perls' Prussian blue staining of aorta sections from 16 weeks old (8 weeks HFD) ApoE$^{-/-}$ mouse (top row) and 26 weeks old (18 weeks HFD) ApoE$^{-/-}$ mouse (bottom row), both injected with HAP-multitag (scale bar is 100 μm).

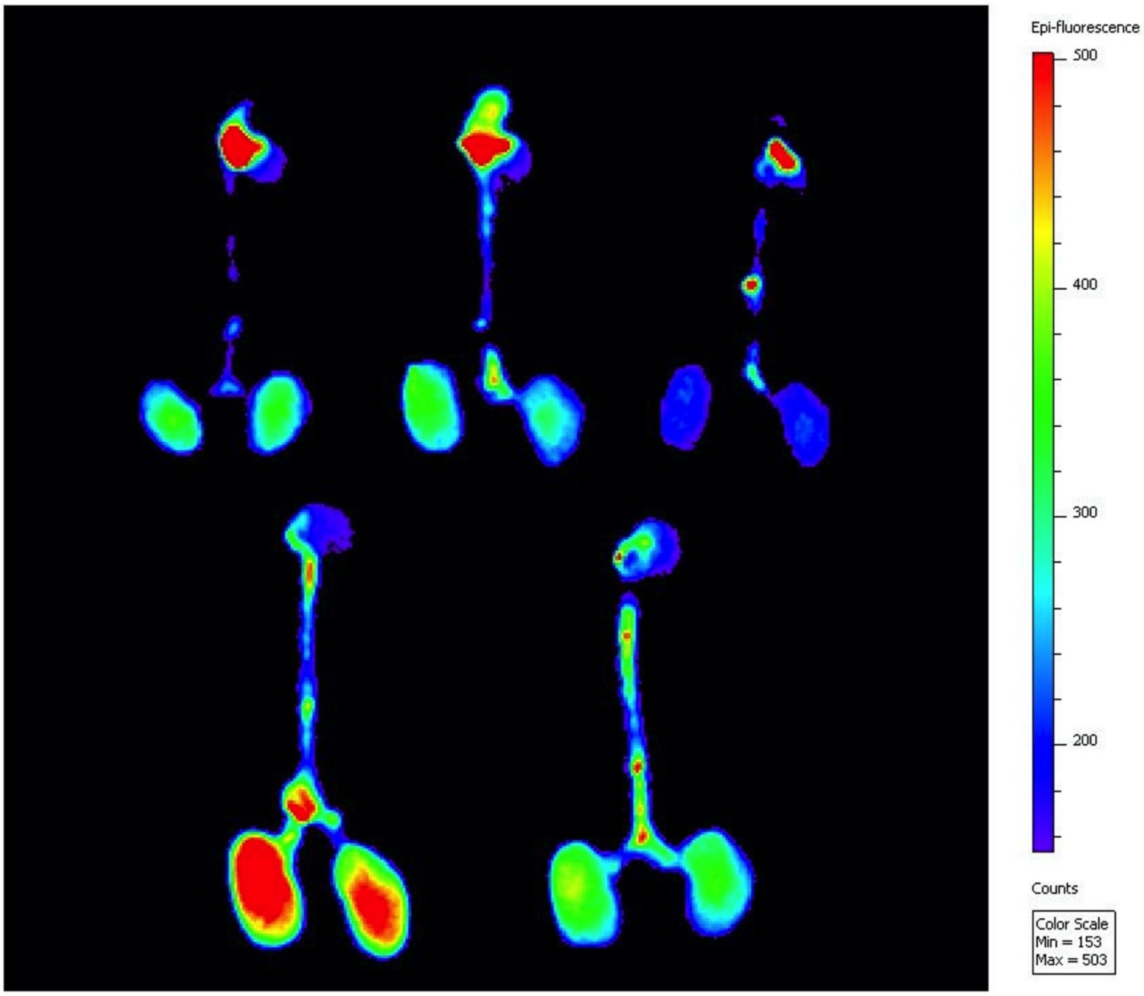


**Figure S4.** *Ex vivo* fluorescence imaging of 5 mice aortas 24 h post *i.v.* injection of OsteoSense® 680EX in 16 weeks old (8 weeks HFD) ApoE$^{-/-}$ mice.

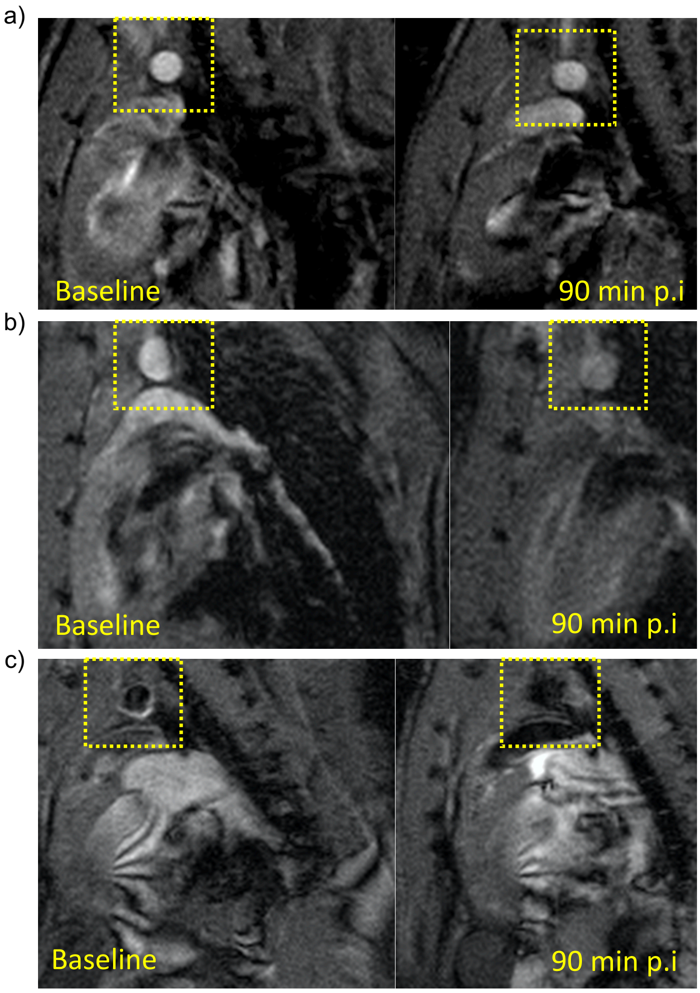


**Figure S5**. $T_1$-weighted MRI before (baseline) and 90 min after *i.v.* injection of $^{68}$Ga-IONP-citrate for a) 12 weeks old (4 weeks HFD) ApoE$^{-/-}$, b) 16 weeks old (8 w HFD) ApoE$^{-/-}$ and c) 26 weeks old (18 weeks HFD) ApoE$^{-/-}$.

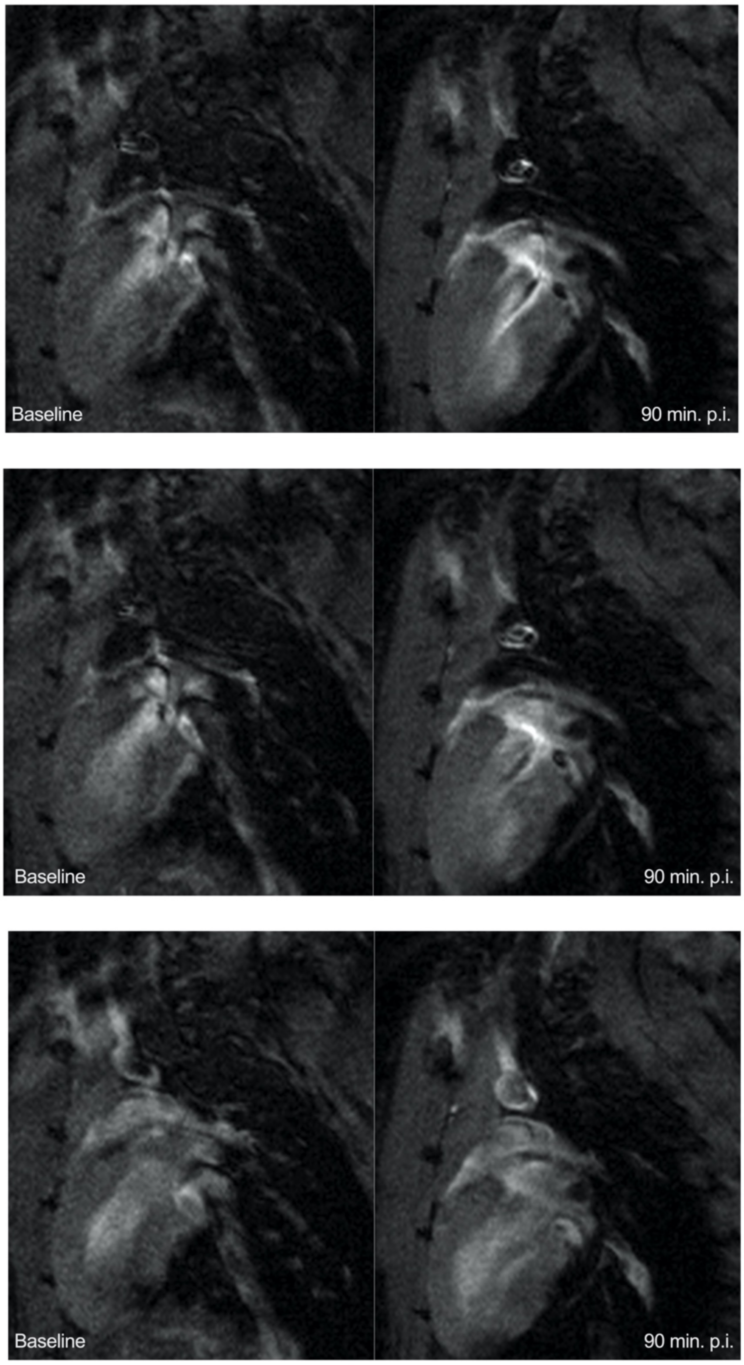


**Figure S6**. $T_1$-weighted MRI before (baseline) and 90 min after *i.v.* injection of HAP-multitag for 16 weeks old (8 w HFD) $ApoE^{-/-}$.
